# Multi-target visual search organisation across the lifespan: Cancellation task performance in a large and demographically stratified sample of healthy adults

**DOI:** 10.1101/307520

**Authors:** Jeroen S. Benjamins, Edwin S. Dalmaijer, Antonia F. Ten Brink, Tanja C.W. Nijboer, Stefan Van der Stigchel

## Abstract

**Highlights:**

- Cancellation tasks are useful clinical tools that probe many cognitive modules
- We used cancellation tests on 523 participants of different ages, sex, and education
- We provide cancellation task norm scores for indices computed from a big sample
- Cancellation indices include attention bias, processing speed and search organisation
- About a quarter of the healthy population shows relatively disorganised search

**Abstract:** It is important that accurate tests exist to assess cognition in various groups of individuals. One popular test of attention and executive functioning is the cancellation task, in which participants perform multi-target visual search to find and ‘cancel’ targets among distractors. Although cancellation tasks have been used extensively with neurological patients, it is only partly clear whether performance is affected by demographic variables such as age and education, which can vary wildly among patients. Here, we describe performance in a sample of 523 healthy participants who participated in a web-based cancellation task. Specifically, we examined indices of spatial bias, processing speed, perseveration and revisiting behaviour, and search organisation. In this sample, age, sex, and level of education did not affect cancellation performance. A cluster analysis identified four cognitive profiles: Participants who make many omissions (N=18), who make many revisits (N=18), who have relatively poor search organisation (N=125), and who have relatively good search organisation (N=362). We advise neurologists and neuropsychologists to exercise caution when interpreting scores pertaining to search organisation in patients: Given the large proportion of healthy individuals with poor search organisation, disorganised search in patients might be pre-existing rather than disorder-related. Finally, we include norm scores for indices of spatial bias, perseverations and revisits, processing speed, and search organisation for a popular cancellation task.

## Introduction

In a cancellation task, participants are required to find and ‘cancel’ (cross out) a number of targets among distractors (Mesulam, 1985). Because targets are spread across the stimulus array, cancellation tasks are sensitive to deficits in spatial attention (Binder, Marshall, Lazar, Benjamin, & Mohr, 1992). As such, they have been a crucial instrument in research on and the diagnosis of visuospatial neglect syndrome, which occurs in 25-50% of stroke patients (Appelros, Karlsson, Seiger, & Nydevik, 2002; Buxbaum et al., 2004; Nijboer, Kollen, & Kwakkel, 2013).

In addition to their diagnostic use in neglect, cancellation tasks have been used to assess cognition in many different groups. For example, ‘invisible’ cancellation tests (in which cancelled targets are not marked) are regularly used to assess short-term memory deficits in stroke patients (Dalmaijer et al., 2018; Husain & Rorden, 2003; Malhotra et al., 2005; Malhotra, Parton, Greenwood, & Husain, 2006; Parton et al., 2006). Another example is search organisation (the extent to which the cancelled targets lie on a sensible path), which has been used as an outcome measure in stroke patients (Dalmaijer et al., 2018; Donnelly et al., 1999; Mark, Woods, Ball, Roth, & Mennenmeier, 2004; Ten Brink et al., 2016; Ten Brink, Van der Stigchel, Visser-Meily, & Nijboer, 2015; Ten Brink, Verwer, Biesbroek, Visser-Meily, & Nijboer, 2017; Woods & Mark, 2007), and children (Woods et al., 2013). For a comprehensive review on what indices can be computed from cancellation tasks, see (Dalmaijer, Van der Stigchel, Nijboer, Cornelissen, & Husain, 2015).

Because of their use in diagnostics and research in populations of a wide variety of backgrounds, it is important that normative data exists for cancellation tasks, and that the effects of demographic factors (e.g. age, sex, and level of education) on cancellation behaviour are known. Although a large body of research (described below) has focussed on demographic effects on traditional cancellation outcome measures (i.e. the number of cancelled targets and task completion times), very little research exists on contemporary measures of interest, such as search organisation. Furthermore, it is unclear how the healthy population performs cancellation tasks.

Here, we describe cancellation behaviour in a large (N>500) sample of healthy adults. We provide norm scores for nearly all currently described outcome measures, and describe demographic effects on these measures. This is of particular import for indices of search organisation, as clinicians and researchers currently have little intuition for what values constitute truly disorganised search. Our norm scores outline where regular behaviour stops, and where irregular behaviour begins. In addition, we employ a data-driven clustering analysis to identify cognitive profiles that exist within the healthy population. The current study and related literature are discussed in greater detail below.

### Known demographic effects on cancellation

Age has been reported to be either a weak (Byrd, Touradji, Tang, & Manly, 2004; Lowery, Ragland, Gur, Gur, & Moberg, 2004) or not a predictor of how long healthy participants take to complete a cancellation task (Brucki & Nitrini, 2008; Saykin et al., 1995). In addition, age did not correlate with spatial bias among healthy adults (Lowery et al., 2004; Saykin et al., 1995).

Years of education did not predict how long healthy adults took to complete a cancellation task (Lowery et al., 2004; Saykin et al., 1995), nor did it predict spatial bias (Lowery et al., 2004; Saykin et al., 1995; Uttl & Pilkenton-Taylor, 2001). However, in a group of people who received very few (under 4) years of education due to their living in secluded communities, there was a difference between literates and illiterates in the number of correctly marked targets found and task completion time (literates found more targets and finished quicker); although within illiterates there was no difference between those that went to school and those that did not (Brucki & Nitrini, 2008).

Gender was not associated with cancellation task duration or spatial bias in some studies (Brucki & Nitrini, 2008; Saykin et al., 1995; Uttl & Pilkenton-Taylor, 2001).

In contrast to the above, Ardila and Rosselli do note “differences appeared for age and schooling” in a letter cancellation task, albeit without further specification on what those differences were (Ardila & Rosselli, 1989). Geldmacher also reported that 30 younger participants (mean age 20.2 years) found more targets and were quicker than 30 older adults (mean age 69.4 years) (Geldmacher, Fritsch, & Riedel, 2000). In addition, Uttl and Pilkenton-Taylor show a minor effect of years of education and of verbal IQ on the number of targets found in 351 healthy adults, but only after detrending their data for the effects of age (Uttl & Pilkenton-Taylor, 2001). Furthermore, Mazaux and colleagues report lower numbers of targets found and slower completion speeds for individuals who were older, of lower education (no schooling or grade school versus high school or university), or female, in a large sample of 1799 healthy participants (Mazaux et al., 1995). Finally, using a 15-minute long letter cancellation task in a sample of 80 healthy participants, Davies and Davies showed that the older (65-72 years) half was slower and less accurate than the younger (18-31 years) half (Davies & Davies, 1975).

Ethnicity has been reported to not correlate with task duration (Lowery et al., 2004; Saykin et al., 1995) or with spatial bias (Lowery et al., 2004; Saykin et al., 1995) in healthy adults. In a study that did show differences in cancellation performance between groups of different ethnicities (that were matched for years of education), no such differences could be observed when statistically controlling for literacy level (Byrd et al., 2004).

In summary, whether age, sex, or education affect the number of cancelled targets or task completion times is unclear, but ethnicity is consistently found to not be a factor. Hence, we focussed on age, sex, and education in the current study.

### Demographic effects on search organisation

The organisation of search paths has traditionally been hard to measure in pen-and-paper cancellation tasks, despite some valiant and creative efforts using for example video recordings or alternating pencil colours (Mark et al., 2004; Samuelsson, Hjelmquist, Jensen, & Blomstrand, 2002; Warren, Moore, & Vogtle, 2008; Weintraub & Mesulam, 1988; Woods & Mark, 2007). However, recent advances in computerised testing have allowed researchers to track when each cancellation was made. As a consequence, search organisation has been an increasingly popular topic of research, and several outcome measures have been proposed (for an overview, see (Dalmaijer et al., 2015)).

For example, it has been shown that disorganised search in stroke patients particularly relates to right hemispheric damage (Ten Brink et al., 2016, 2015; Weintraub & Mesulam, 1988), and some studies (Rabuffetti et al., 2012; Samuelsson et al., 2002; Ten Brink et al., 2015) but not others (Mark et al., 2004) relate it to neglect. Search organisation in neglect patients has also been shown to be unaffected by noradrenergic medication (Dalmaijer et al., 2018). In addition, revisiting behaviour (i.e. re-marking previously cancelled targets) has also been related with neglect (Na et al., 1999; Nys, Van Zandvoort, Van der Worp, Kappelle, & De Haan, 2006; Rusconi, Maravita, Bottini, & Vallar, 2002).

Importantly, deficiencies in search organisation could reveal more subtle cognitive problems compared to traditional outcome measures such as the number of omissions, as these are thought to be less sensitive to compensatory strategies. However, this requires that clinicians and researchers can accurately interpret individual patients’ search organisation scores, which is currently very challenging due to a lack of normative data.

### Current study

For the current study, we investigated multi-target visual search in a sample of 523 healthy participants who took part in a web-based cancellation task. This sample was stratified for age and level of education, in order to allow for statistically meaningful analyses of demographic factors on cancellation measures. Level of education was measured in six categories of formal qualifications: None (e.g. for people who only completed primary school), secondary school (GCSE), College (A levels), university undergraduate (e.g. BSc or BA), university graduate (e.g. MPhil, MSc, or MA), and university or medical doctorate (e.g. DPhil, PhD, or MD).

Typically, how well participants perform on a cancellation task depends on four cognitive modules: Spatial bias, short-term memory, processing speed, and search organisation (often considered to reflect executive function). We assessed the demographical effects on quantifications of these modules. Furthermore, our sample is large enough to allow for the employment of ‘Big Data’ tools to find statistical regularities in datasets. Specifically, we performed a cluster analysis that identified participants with similar cognitive profiles in a data-driven way, using quantifications of the four cognitive modules outlined before.

## Methods

### Procedure

Participants were recruited through Prolific Academic, a website that allows for the recruitment of participants for web-based questionnaires and tasks. We attempted to select a stratified sample of participants by creating 30 different Prolific Academic links (5 age groups x 6 education levels) that were open to 20 participants each. Each of these linked to the same cancellation task (described below). All Prolific Academic links were kept active for six days including a weekend, after which they were deactivated, and participants who completed the task received their reward through Prolific Academic.

In total, 535 individuals took part in our experiment (described in **Table 1**). Out of those, 523 participants cancelled one or more targets, and were included in our analyses.

**Table 1.** Descriptives statistics for the sample of healthy volunteers that took part in our online cancellation task. Age is indicated in years.

| **Education** | **N** | **% male** | **Mean age** | **Age range** |
| --- | --- | --- | --- | --- |
| **No formal qualifications** | 28 | 54 | 39 | 20 - 79 |
| **Secondary school**<br>(GCSE) | 116 | 44 | 42 | 19 - 84 |
| **College**<br>(A levels) | 101 | 47 | 43 | 19 - 73 |
| **Undergraduate degree**<br>(e.g. BA, BSc) | 124 | 45 | 44 | 23 - 75 |
| **Graduate degree**<br>(e.g. MA, MSc, MPhil) | 99 | 63 | 44 | 22 - 79 |
| **Doctorate degree**<br>(e.g. DPhil, PhD, MD) | 67 | 57 | 42 | 23 - 73 |
| **Total** | 535 | 50 | 43 | 19 - 84 |

### Cancellation Task

Participants were presented with a Landolt C cancellation task (Parton et al., 2006) with 64 targets that were circles with an opening on top, and 128 distractors that were circles with an opening at the bottom and circles without an opening (**Figure 1**). They were instructed to click on all the targets while ignoring the distractors. When a target was clicked, a cross appeared to mark its cancellation. There was no time limit, participants could click a button to end the task when they thought they had found all targets. The search array was generated using CancellationTools (Dalmaijer et al., 2015), and the task was programmed in JavaScript and implemented on the LimeSurvey platform (version 2.05+ build 141229) available to author JB.

**Figure 1.**
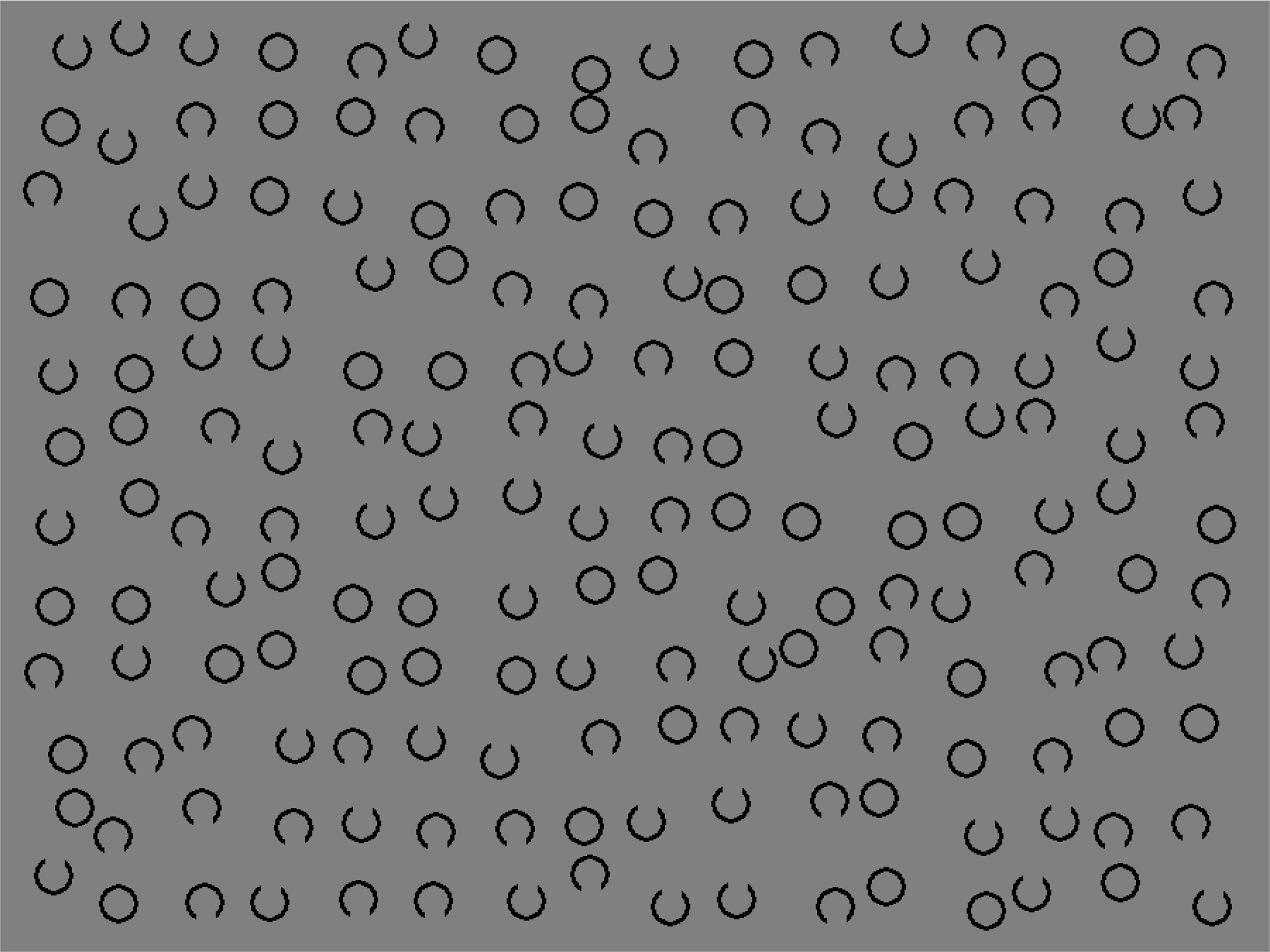
Landolt C cancellation task with 64 targets (circles with a top opening) and 128 distractors (circles with a bottom opening, and without an opening).

To minimize data loss due to screen resolution limitations of the devices on which participants performed the task, they could only continue if their screen was large enough. This was enforced by presenting a neutral image (a picture of a grain field) and a button in participants’ browsers, and only allowing them to continue when they could see both without scrolling. The task could only be accessed on desktop computers and laptops.

### Outcome measures

Data from 523 participants were first processed using CancellationTools (Dalmaijer et al., 2015), which uses task target coordinates and participants’ click coordinates and timestamps to compute several indices that relate to cancellation performance.

Cancellation indices that relate to attentional bias include the number of omissions (non-cancelled targets), the difference between the number of omissions on the left and right halves of the search array, and the centre of cancellation (the average horizontal cancellation location when the left-most target location was defined as −1 and the right-most as 1; introduced by (Binder et al., 1992)). Delayed revisits (when a participant re-cancelled a previously cancelled target after cancelling different targets) correlate with short-term memory, particularly when cancellations are unmarked (Dalmaijer et al., 2018; Malhotra et al., 2005, 2006; Parton et al., 2006). Immediate revisits (sometimes referred to as ‘perseverations’) are repeated clicks on the last cancelled target, and are thought to reflect impulse control or motor issues. Because participants were asked to stop the task after they thought they had found all targets, the total task duration can be used as an index of processing speed, as can the time between consecutive cancellations. Finally, search organisation can be quantified as the distance between consecutive cancellations (with lower distances reflecting better search organisation), by using the standardised angle (a number that is closer to 1 for cardinal, and closer to 0 for diagonal inter-cancellation angles), the highest correlation between cancellation rank number and horizontal or vertical cancellation location (best R; (Mark et al., 2004; Woods & Mark, 2007)), and the number of times a cancellation path crosses divided by the total number of cancellations (intersection rate). These indices and their origins are reviewed by (Dalmaijer et al., 2015).

As described in the introduction, cancellation task performance depends primarily on spatial bias, short-term memory, processing speed, and search organisation. Each of these can be quantified using a number of metrics describe above, and here we chose to focus on the total number of omissions to quantify spatial bias, the total number of revisits (immediate and delayed) to quantify short-term memory, the task duration to quantify processing speed, and the best R to quantify search organisation. Although more could have been included, the four selected outcome measures are representative of the outlined cognitive modules (see the correlation matrix plotted in **Figure 2**). Including additional measures would have blunted our statistical tools by requiring a more stringent correction for multiple comparisons, and by adding additional dimensions to the data matrix that was fed into a cluster analysis. By choosing four representative measures, we aimed to avoid the '*curse of dimensionality*': The tendency of clustering algorithms to perform increasingly poorly when including more features (Bellman, 1957). Note that for transparency and usability, we included all eleven computed measures in reported norm scores (**Table 2**) and correlations (**Figure 2**), as these constitute descriptive statistics rather than tests.

**Table 2.** Percentile scores at 10-point intervals, starting from the 5^th^ and ending at the 95^th^ percentile. Where possible, values are listed from bad performance to good performance, so that the 1^st^ percentile is the worst performance, and the 100^th^ the best. The one exception to this is the omission bias, which is listed from lowest to highest values, from leftward bias (more omissions on the right) to rightward bias (more omissions on the left). Note that ‘immediate revisits’ are sometimes referred to as ‘perseverations’ in the literature.

|  | Percentile |  |  |  |  |  |  |  |  |  |
| --- | --- | --- | --- | --- | --- | --- | --- | --- | --- | --- |
|  | 5 | 15 | 25 | 35 | 45 | 55 | 65 | 75 | 85 | 95 |
| **Omissions** | 10 | 5 | 3 | 2 | 1 | 1 | 0 | 0 | 0 | 0 |
| **Omission bias (right - left)** | -2 | -1 | 0 | 0 | 0 | 0 | 0 | 1 | 1 | 3 |
| **Centre of cancellation** | 0.10 | 0.09 | 0.08 | 0.08 | 0.08 | 0.08 | 0.08 | 0.07 | 0.06 | 0.04 |
| **Immediate revisits** | 5 | 2 | 1 | 0 | 0 | 0 | 0 | 0 | 0 | 0 |
| **Delayed revisits** | 2 | 1 | 0 | 0 | 0 | 0 | 0 | 0 | 0 | 0 |
| **Duration (sec)** | 132 | 108 | 99 | 91 | 85 | 79 | 73 | 69 | 65 | 55 |
| **Inter-cancel time (sec)** | 2.18 | 1.71 | 1.57 | 1.45 | 1.35 | 1.25 | 1.16 | 1.11 | 1.01 | 0.88 |
| **Inter-cancel distance (pix)** | 197 | 163 | 155 | 149 | 144 | 137 | 132 | 126 | 119 | 106 |
| **Standardised angle** | 0.53 | 0.56 | 0.59 | 0.61 | 0.63 | 0.66 | 0.70 | 0.75 | 0.78 | 0.81 |
| **Best R** | 0.40 | 0.57 | 0.68 | 0.80 | 0.89 | 0.94 | 0.96 | 0.98 | 0.99 | 0.99 |
| **Intersections rate** | 0.34 | 0.22 | 0.16 | 0.11 | 0.08 | 0.05 | 0.03 | 0.02 | 0.00 | 0.00 |

**Figure 2.**
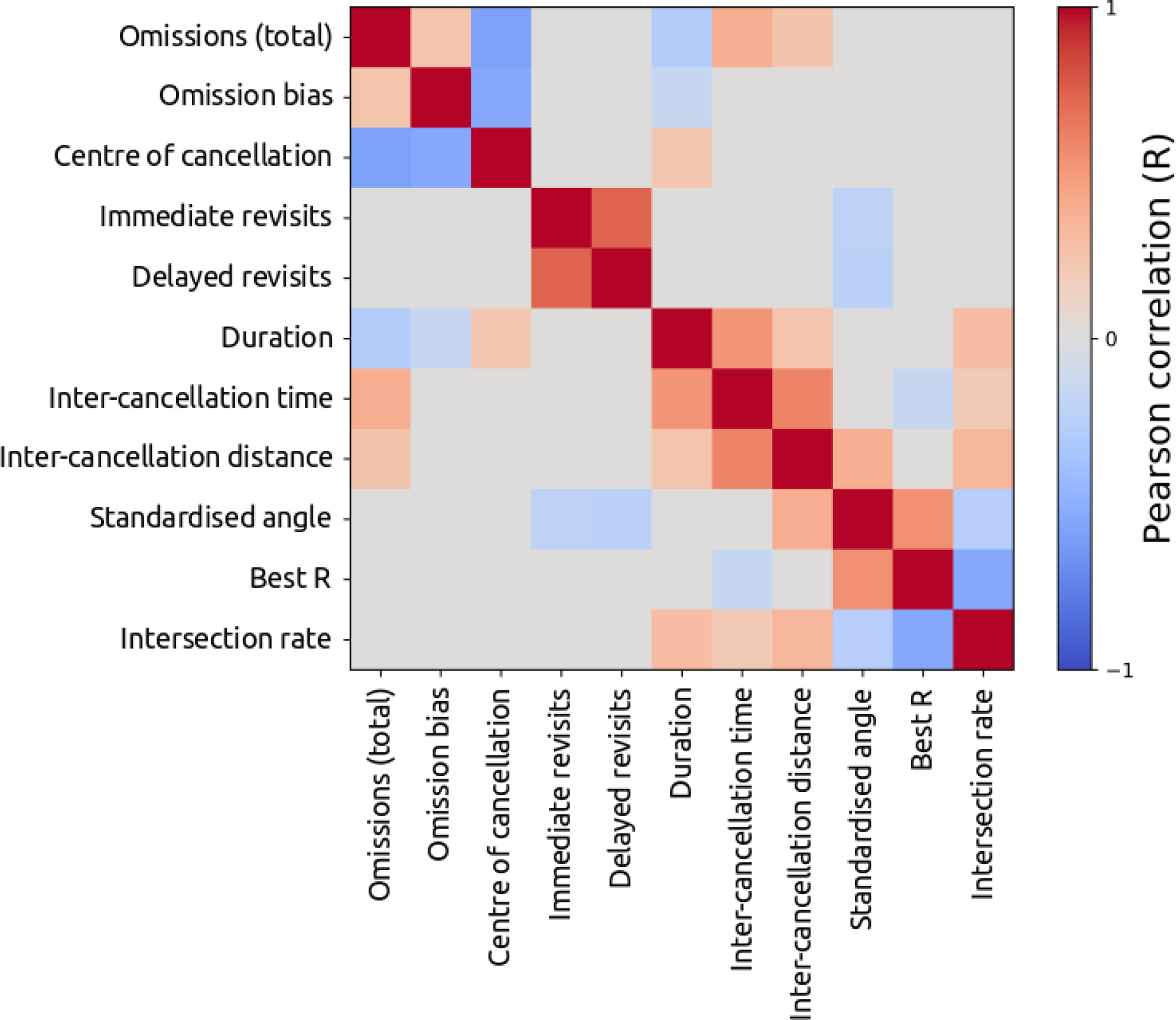
Correlation matrix of the most important measures computed by CancellationTools, given the data collected from our sample. Only R values that were significant at a Bonferroni-corrected alpha of 0.05 are visualised. From top to bottom on the y-axis (and left to right on the x-axis): number of omissions, omissions bias (omissions on right half minus those on the left half), horizontal centre of cancellation, number of immediate revisits (sometimes referred to as ‘perseveration’), number of delayed revisits, task duration (i.e. time until participant stopped the task), average time between consecutive cancellations (inter-cancellation time), average distance between consecutive cancellations (mean inter-cancellation distance), standardised angle between consecutive cancellations, best R, and rate of intersections.

It should be noted that the number of omissions is not the most elegant measure of spatial bias in neglect patients. In healthy controls, the number of omissions is also sensitive to participants who misunderstood the task or were unmotivated to do it (and therefore skipped many targets), whereas bias measures show very little variability (**Table 2**). Hence, it is a pragmatic choice for the analyses outlined below. For interpretation of patient scores, norm values are provided for the left-right omission bias and centre-of-cancellation measures in **Table 2**.

In addition, it should be noted that the cancellations were marked in our experiment, and that revisits are thus more likely to reflect problems with inhibition or task understanding than with short-term memory.

### Statistical tests

To investigate the effect of age on cancellation performance, we computed the Pearson correlation coefficient for age and each of the four selected measures: number of omissions, number of revisits, task duration, and best R. We did this within the entire sample, and within each type of education. To account for multiple comparisons, a Holm-Bonferroni correction was employed (Holm, 1979).

To investigate the effect of sex and education type on cancellation performance, we performed ANCOVAs with age as covariate, and sex and education as factors. Sex had two levels (male and female), and education had 6 levels: None (primary school only), secondary, college (A levels), university undergraduate (e.g. BSc, BA), university graduate (e.g. MSc, MA, MPhil), university or medical doctorate (DPhil, PhD, MD). One ANCOVA was performed for each of the four selected cancellation measures: number of omissions, number of revisits, task duration, and best R.

### Cluster analysis

To investigate whether distinct sub-groups exist in the healthy population, we applied k-means clustering on aforementioned measures (number of omissions, number of revisits, best R, and task duration). This type of analysis provides a data-driven way to identify clusters of participants who share behavioural traits (and thus potentially cognitive traits), which is important to consider when observing patients whose cognitive background is unknown. Post-clustering chi-square tests allowed for comparison of demographic factors between the clusters.

The computed metrics are in non-unified spaces. For example, best R scores exist in a range between 0 and 1, but the duration is measured in seconds (and can thus range from under ten to several hundreds). These ranges are relatively arbitrary, but will bias clustering algorithms nonetheless. To account for this, values are min-max transformed within each measure to a range between 0 and 1.

These transformed values were then fed into a series of k-means cluster analyses (for a historical overview of the algorithm, see (Jain, 2010)), as a matrix of 523 observations by 4 features. K-means clustering requires a user to set the number of clusters k, and then organises k cluster centroids in a random order. On each iteration of the algorithm each point is assigned to the closest cluster centroid (samples are thus members of only one cluster), and new cluster centroids are computed as the average location of all samples assigned to a cluster. The algorithm converges when no the cluster centres do not change between iterations any further.

After convergence, each sample can be assigned a silhouette value (Rousseeuw, 1987). The silhouette value of sample *i* is computed by subtracting the average distance between sample *i* and all samples in its cluster from the average distance between sample *i* and all samples in the nearest other cluster, and by dividing this by which of those two values was the highest. The resulting value can range from −1 to 1. A value of 1 means that sample *i* is perfectly aligned with its assigned cluster. Conversely, a value of −1 indicates that sample *i* is perfectly aligned with a cluster it was not assigned to. A value of 0 indicates that sample *i* lies in between its assigned cluster and a different cluster. All silhouette values can be averaged to compute the *cluster coefficient*. A rule of thumb is that values between 0 and 0.25 indicate that the data shows no cluster structure, values between 0.25 and 0.5 indicate potentially artificial clustering, values between 0.5 and 0.70 indicate reasonable clustering, and values between 0.70 and 1 indicate good clustering (Kaufman & Rousseeuw, 1990).

A common way of identifying how many clusters exist in a dataset is to cycle through several k-means clustering analyses, and choose the number of clusters (k) for which the cluster coefficient is highest. Here, 10 cluster analyses (k=1 to k=10) will be compared, and the best solution will be chosen based on the cluster-coefficient. A coefficient of over 0.5 will be considered evidence for a reliable cluster structure being present in the data.

For visualisation purposes, the data are subjected to a multi-dimensional scaling procedure. More specifically, the data is processed using t-distributed stochastic neighbour embedding (t-SNE; (Van der Maaten & Hinton, 2008)). This technique aims to reduce the number of dimensions in which the data is defined (typically to two), which allows for easier plotting within a single graph. It does so by preserving local but not necessarily global structure, making distances in the resulting plot relatively arbitrary. It is important to note that this is a stochastic process, and will thus not always produce the same result. It is also important to note that t-SNE is only used here to produce illustrations; not as a pre-processing step before clustering, and not for the purpose of making statistical comparisons.

## Results

### Normative data for cancellation measures

Data for nearly all cancellation measures that CancellationTools computes have been summarised in **Table 2**. These values correspond with the 5^th^, 15^th^, 25^th^, 35^th^, 45^th^, 55^th^, 65^th^, 75^th^, 85^th^, and 95^th^ percentile, and can be used as normative data. Note that the values are listed according to performance, not according to value. For example, for the best R higher values indicate better performance, but for the number of omissions lower values indicate better performance.

### Relations between cancellation measures

Before analysing selective effects of demographics on cancellation measures, it is useful to understand how measures relate to each other. We computed the Pearson correlations between the values for each individual for each measure computed by CancellationTools. The resulting correlation matrix is plotted in **Figure 2**. Qualitatively, four groups of measures appear to exist. The first group relates to the number and bias of omissions, and includes the total number of omissions, the omission bias (omissions on the right half minus omissions on the left half), and the horizontal centre of cancellation (introduced by (Binder et al., 1992)); which all correlate strongly with each other, and to a lesser extent with duration measures. The second group of measures relates to revisits, and shows that immediate and delayed revisits correlate strongly with each other, but not with any other measures (save from the standardised angle, a metric of search organisation introduced by (Dalmaijer et al., 2015)). The third group relates to timing, and includes the task duration, the average time between consecutive cancellations; both correlate with each other, as well as with the average distance between consecutive cancellations and the total number of omitted targets. The final group of measures relates to search organisation, and includes the average distance between consecutive cancellations, the standardised angle (high for better search organisation, introduced by (Dalmaijer et al., 2015)), the best R (high for better search organisation, introduced by (Mark et al., 2004)), and the rate of cancellation path intersections (low for better search organisation, introduced by (Donnelly et al., 1999)).

As is evident from the correlation matrix (**Figure 2**), the most representative measures for the aforementioned groups are the total number of target omissions, the total (immediate and delayed) number of target revisits after initial cancellation, the task duration, and the best R.

### Effect of age on cancellation

Age is not a significant predictor of any selected measure, save from the duration (time-on-task) for which it accounted for 10 percent of the variance in the whole sample (**Table 3**). On average, older participants required more time to complete the task, at a rate of 0.59 seconds of time-on-task per year-of-age.

**Table 3.** Explained variance (R^2^) for cancellation measures when predicted with age in a linear regression, either in the entire sample (“All” column), or within a group with the same educational background. Bold R^2^ values indicate the associated p value was below 0.05 (exact p values are reported within the same cell). Asterisks indicate statistical significance after Holm-Bonferroni correction for multiple comparisons.

|  | **All**<br>(N=523) | **None**<br>(N=27) | **Secondary**<br>(N=112) | **College**<br>(N=100) | **Undergrad**<br>(N=124) | **Grad**<br>(N=94) | **Doctorate**<br>(N=66) |
| --- | --- | --- | --- | --- | --- | --- | --- |
| **Omissions** | 0.0000011 | 0.054 | 0.00039 | 0.033 | 0.00074 | 0.0016 | 0.0040 |
| **Revisits** | 0.00039 | 0.0081 | 0.020 | 0.00049 | 0.0069 | 0.023 | 0.049 |
| **Best R** | 0.0026 | 0.047 | **0.037**<br>$p=0.0418$ | 0.022 | 0.0029 | 0.022 | 0.0027 |
| **Duration** | **0.10\***<br>$p<0.00001$ | 0.064 | **0.29\***<br>$p<0.00001$ | **0.080**<br>$p=0.00459$ | **0.13\***<br>$p<0.0001$ | 0.036 | **0.12**<br>$p=0.00421$ |

### Effects of sex and education on cancellation

An ANCOVA with age as co-variate, and sex (2 levels: male and female) and education (6 levels: none, secondary, college, undergrad, grad, doctorate) as fixed factors revealed a statistically (*p* = 0.026) but not practically (η^2^ = 0.01) significant effect of sex on the **number of omissions** F(1, 510) = 5.00, *p* = 0.026, η^2^ = 0.01; no effect of level of education, F(5, 510) = 1.00, *p* = 0.420; no sex*education interaction effect, F(5, 510) = 1.08, *p* = 0.371; and no effect of age, F(1, 516) = 0.01, *p* = 0.936.

An ANCOVA with age as co-variate, and sex (2 levels: male and female) and education (6 levels: none, secondary, college, undergrad, grad, doctorate) as fixed factors revealed no effect of sex on the **total number of revisits**, F(1, 510) = 1.21, *p* = 0.272; no effect of level of education, F(5, 510) = 1.29, *p* = 0.266; no sex*education interaction effect, F(1, 510) = 1.75, *p* = 0.122; and no effect of age, F(1, 510) = 0.27, *p* = 0.607.

An ANCOVA with age as co-variate, and sex (2 levels: male and female) and education (6 levels: none, secondary, college, undergrad, grad, doctorate) as fixed factors revealed no effect of sex on **best R**, F(1, 510) = 0.55, *p* = 0.458; no effect of level of education, F(5, 510) = 1.22, *p* = 0.298; no sex*education interaction effect, F(1, 510) = 0.29, *p* = 0.916; and no effect of age, F(1, 510) = 2.06, *p* = 0.152.

An ANCOVA with age as co-variate, and sex (2 levels: male and female) and education (6 levels: none, secondary, college, undergrad, grad, doctorate) as fixed factors revealed no effect of sex on **task duration**, F(1, 510) = 2.51, *p* = 0.114; no effect of level of education, F(5, 510) = 1.38, *p* = 0.229; no sex*education interaction effect, F(1, 510) = 0.94, *p* = 0.452; and a statistically significant effect of age, F(1, 510) = 58.00, *p* < 0.001, η^2^ = 0.10.

### K-Means Clustering

The solution of a 4-means cluster analysis proved to produce the highest cluster-coefficient (**Figure 3A**), and should thus be taken as the best solution. The value is over 0.5, and should thus be treated as evidence for a reliably clustered (multi-modal) structure being present in the data (for a full silhouette plot, see **Figure 3B**).

**Figure 3.**
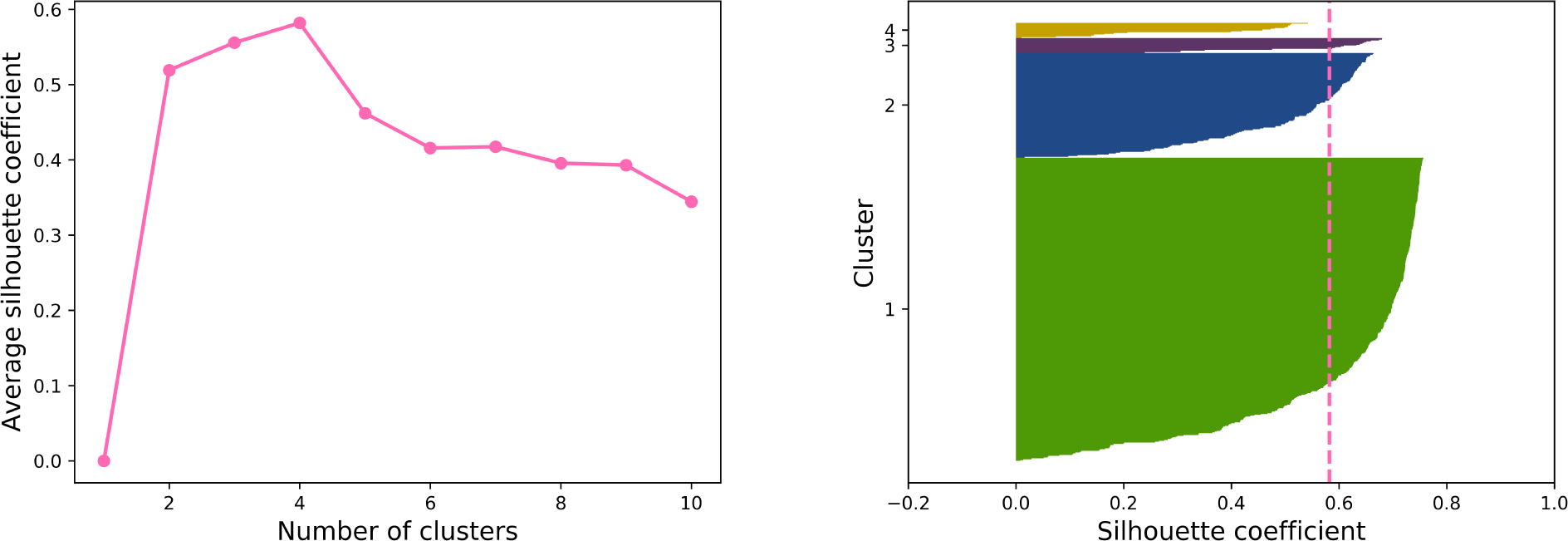
**A)** Cluster coefficient (y-axis) as a function of cluster size k (x-axis). A higher cluster coefficient indicates better clustering. The best solution here is for k=4 clusters. **B)** Silhouette plot of the k=4 cluster solution. Each sample is organised along the y-axis, sorted from high to low cluster coefficient (x-axis) within each cluster. The dotted line (pink) represents the average cluster coefficient of the k=4 solution.

In the best solution (k=4), the first cluster consisted of 18 participants who were best characterised by their high number of omissions, the second cluster consisted of 18 participants who were best characterised by their high number of revisits, the third cluster consisted of 125 participants who were best characterised by their low best R values, and the fourth cluster consisted of 362 participants who were best characterised by their high best R values. Task duration was slightly lower in the high omission group, and was highly similar between the other clusters (**Figure 4A**). Feature space was plotted in **Figures 4A and 4B**, and reduced space (using t-SNE) in **Figure 4C**. Each cluster’s average scores on the four features (best R, task duration, number of revisits, and number of omissions) were plotted in **Figure 5A**. It should be noted that the values in these figures are min-max scaled to fit between 0 and 1 (**Figure 4A-B and 5A**), or warped into arbitrary space (**Figure 4C**). Non-transformed values are listed in Table 1.

**Figure 4.**
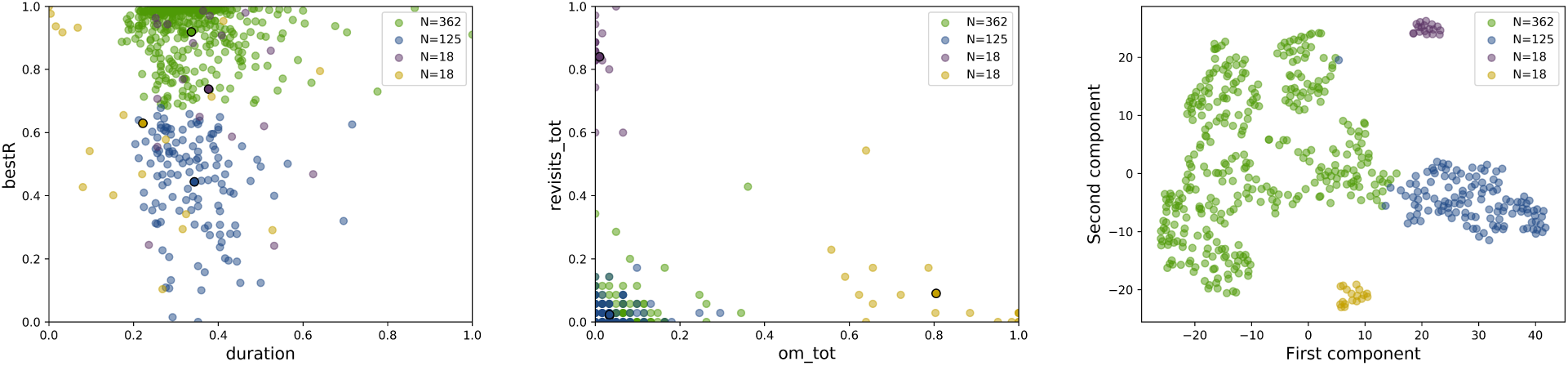
Descriptive plots of the four clusters, one with structured search organisation (N=362, green), one with unstructured search organisation (N=125, dark blue), one with a high number of revisits (N=18, purple), and one with a high number of omissions (N=18, yellow). Each dot in the scatter plots represents an individual participant. Individual samples are indicated as circles. Cluster centroids are indicated by circles with black outlines. **A)** Scatter plot of best R (y-axis) and duration (x-axis), clearly dissociating the clusters with organised (N=362, green) and disorganised search (N=125, dark blue). Values are min-max scaled to fit between 0 and 1. **B)** Scatter plot of the number of revisits (y-axis) and the number of omissions (x-axis), clearly dissociating between the high revisits cluster (N=18, purple), the high omissions cluster (N=18, yellow), and the other two clusters. **C)** Scatter plot in along two components. Space was reduced from four features to two using t-distributed stochastic neighbour embedding (t-SNE). In this reduced dimensional view, it is easier to see the four separate clusters.

**Figure 5.**
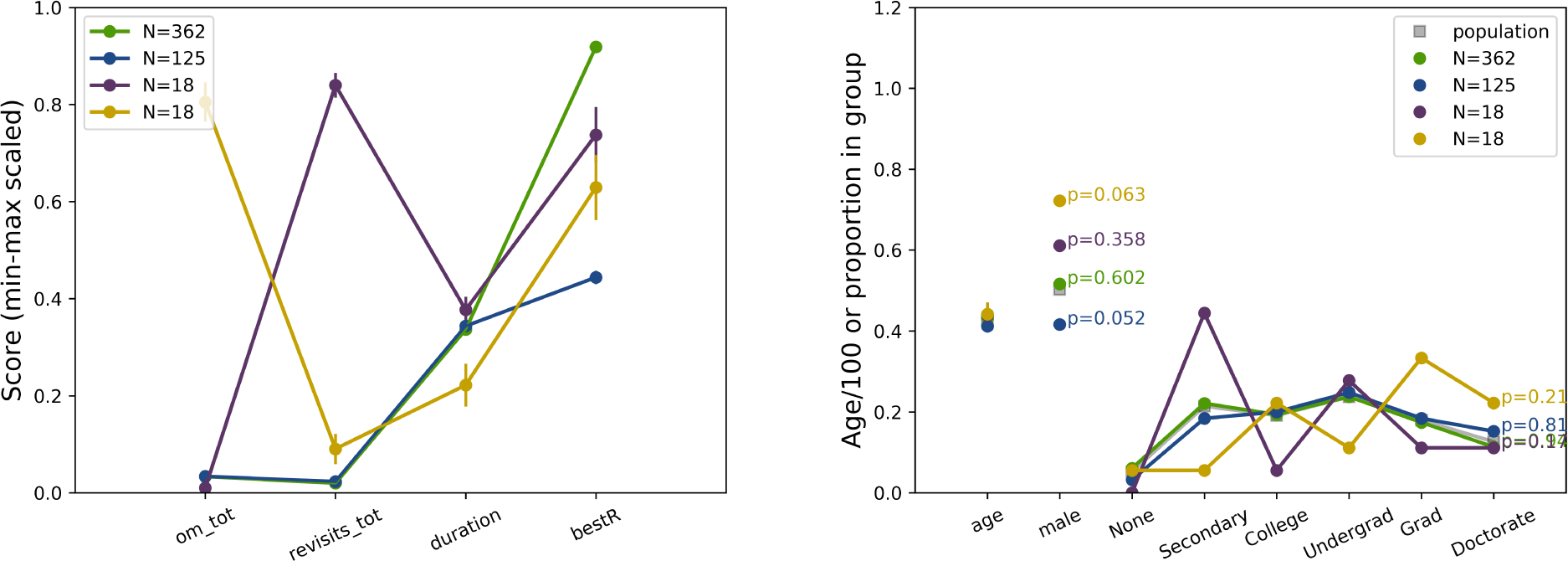
Descriptive plots of the four clusters, one with structured search organisation (N=362, green), one with unstructured search organisation (N=125, dark blue), one with a high number of revisits (N=18, purple), and one with a high number of omissions (N=18, yellow). Solid lines represent within-cluster averages, and error bars the standard error of the mean. **A)** Min-max scaled averages (y-axis) for four features (number of omissions, number of revisits, task duration, and best R. The clusters differ from each other on these features per definition, as these are the values used to identify them. **B)** Average age, proportion of males, and proportions of highest education (None, secondary school, college, university undergraduate, university graduate, and university or medical doctorate) in all four clusters. Annotated p-values report significance of chi-square tests that tested whether a cluster’s make-up significantly differed from the whole sample (plotted in grey squares).

None of the clusters differ significantly from the whole sample in their proportion of males (all p > 0.052 in one chi-square test per cluster), or in their proportion of different levels of education (all p > 0.17 in one chi-square test per cluster); see **Figure 5B**. Average age also does not differ between the clusters (**Figure 5B**). These results indicate that cancellation performance does not predict demographic values, which mirrors the aforementioned results that showed no evidence that demographics predict cancellation performance (save from a minor effect of age on duration).

## Discussion

We describe multi-target visual search performance in a sample of 523 healthy individuals who participated in a web-based cancellation task. In this sample, age, sex, and level of education did not affect cancellation performance. The only exception to this was age predicting 10 percent of the variance in task duration. Cluster analyses identified four cognitive profiles. They can be best described as individuals who make many (around 10 or more) omissions (N=18), individuals who make many (around 7 or more) revisits (N=18), individuals with relatively bad search organisation (best R < 0.7; N=125), and individuals with relatively good search organisation (N=362). Note that the cut-off scores were not pre-defined, but rather derived from the (data-driven) clustering analysis, and are thus of a descriptive nature. Importantly, task duration was not different between the groups with better and worse search organisation, which suggests search organisation is a trait that does not necessarily impact performance.

We conclude that cancellation task outcome measures are not affected by age, sex, or education. In addition, we conclude that a significant part of the healthy population shows relatively poor search organisation. We provide norm scores (**Table 2**) based on the data from our healthy participants to help interpret cancellation performance in patients.

### Demographic effects on cancellation

Previous research has primarily focussed on years of education rather than type of education (Lowery et al., 2004; Saykin et al., 1995; Uttl & Pilkenton-Taylor, 2001), or only used very coarse differences in type of education such as “no school or grade school” versus “high school or university” (Mazaux et al., 1995). In addition, some studies only found effects of years of education on cancellation performance when statistically controlling for age (Uttl & Pilkenton-Taylor, 2001).

Here, we tested individuals from a wide variety of educational backgrounds: no further schooling (primary school), secondary school, college (A levels), university undergraduate (BSc or BA), university graduate (MSc, MA, MPhil), and university or medical doctorate (DPhil, PhD, MD). In addition, we controlled for age when statistically investigating the effects of education.

Our results show that age, sex, and level of education did not affect cancellation outcome measures, with the exception of age accounting for a small part (ten percent) of the variance in cancellation task completion times. These results are in line with existing research (Lowery et al., 2004; Mazaux et al., 1995; Saykin et al., 1995). Despite its effect on task duration, age did not affect omissions, spatial bias, revisits, or search organisation.

### Cancellation in healthy individuals

The current results demonstrate that cancellation tasks can be used to assess individual differences in some aspects of cognition. Four different groups emerged from a k-means cluster analysis that was run on 523 observations with 4 features each. The chosen features were the number of omitted targets, the number of revisits (delayed and immediate combined), the best R to index search organisation, and the task duration. These were chosen to be the most representative of the cognitive domains that a cancellation task aims to test: spatial attention (omissions), short-term memory (revisits), search organisation / executive functioning (best R), and processing speed (time-on-task).

The resulting clusters included two small clusters, with 18 members each. The first of these was a cluster with many omissions. This cluster did not show a different spatial bias from other clusters, and thus might instead just reflect a group of participants that were not motivated to perform the task in the correct way. This was supported by the cluster’s comparatively low time-on-task.

The second small (18 members) cluster showed many revisits. This is surprising behaviour, as markings were visible during the task. Hence, revisits here might not necessarily reflect short-term memory, but rather impulse control, or simply a misunderstanding of the task.

A larger cluster with 125 members was characterised by relatively poor search organisation. Within this group, there was still a reasonable variability of best R scores, and of task durations (time-on-task). This demonstrated that although these participants share a cognitive profile, there are still individual differences.

The final cluster identified here consisted of 362 members, and was characterised by well-structured search, but this group did not differ from the group with poor search organisation with respect to task duration.

In a large proportion of the cluster with good search organisation, best R scores plateaued, although some variability remained. Perhaps more interesting is that there was still considerable variability in the time-on-task even among participants with plateau-level search organisation. This, and the fact that the poor and good search clusters did not differ in average task duration, suggests that processing speed is an individual characteristic that is independent from the ability to search in an organised way.

### Interpretation of cancellation measures

The results from a healthy sample presented here outline what one could learn from a stroke patient’s test scores on a cancellation task. In particular, it illustrates that scores related to spatial bias (target omissions, left-right omission difference, and centre of cancellation) are reasonably low in healthy participants, and thus that even small deviations could be indicative of potential neglect. On the other hand, a low score on a search organisation measure does not necessarily imply that a patient’s stroke is to blame. As demonstrated here, a large proportion of the healthy population will actually score relatively poorly, and thus a poor score after stroke is not necessarily a product of that stroke. Unless a pre-stroke measurement exist, low search organisation scores should be interpreted with respect to normative data. In other words, we would argue that only highly disorganized search is indicative of cognitive problems, as some variation in search organisation naturally exists.

More specific direction is given in **Table 2**, where normative data is provided on the basis of all 523 healthy participants included in our analysis. It should be noted that these norm scores are specific to the Landolt C cancellation task (**Figure 1**) used here (and by (Dalmaijer et al., 2018; Malhotra et al., 2006; Parton et al., 2006)), and that using other cancellation tasks might yield different results. Many factors influence cancellation behaviour, including the target to distractor ratio (Geldmacher, 1996), similarity of targets and distractors (Geldmacher, 1998), the surface area and density of the cancellation task (Geldmacher et al., 2000), the placement of stimuli on a strict grid (Hills & Geldmacher, 1998), and whether or not cancellations were marked (Parton et al., 2006). This literature suggests a need for normative data for each unique cancellation task (e.g. Bell’s test, Star Cancellation, Balloons Test). In addition, not all individuals (patient or healthy) necessarily share the same underlying mechanism for disorganised search, and more research is needed to fully understand what drives search organisation. Finally, our participants were not instructed to search in an organised fashion (whereas patients who sometimes feedback during the task), completed only one cancellation task (unlike patients who are regularly tested), and did the task without any consequences of the outcome (whereas test outcomes contribute to patients’ diagnoses and treatment). Our healthy participants seemed well motivated, given their high accuracy (over 85% made fewer than 5 omissions) and good completion times (1-2 minutes), as reported in Table 2. In future studies, it could be informative to evaluate whether explicit instructions affect search organisation in patients and healthy individuals, and whether repeated testing differentially alters search behaviour between the two groups.

Other researchers have noted that hospitalised elderly patients often perform worse on a large battery of cognitive tests than younger hospitalised patients, and compared to healthy peers; even if no brain injury is present (Woods, Mark, Pitts, & Mennemeier, 2011). This highlights that brain injury is not the only cause of cognitive impairment, and underscores a need for broad cognitive assessment of (elderly) patients with and without brain injury. Cancellation tasks are easy, short, and inexpensive tests that probe a wide body of independent cognitive modules. With the data and norm scores provided here, clinicians and researchers can employ cancellation tests with their patients, and interpret the output with a reasonable degree of confidence.

## Acknowledgements

We would like to thank Masud Husain for reading and commenting on an earlier version of this manuscript. ESD was supported through a European Union FP7 Marie Curie ITN grant (606901).

## References

Appelros, P., Karlsson, G. M., Seiger, A., & Nydevik, I. (2002). Neglect and Anosognosia After First-Ever Stroke: Incidence and Relationship to Disability. Journal of Rehabilitation Medicine, 34(5), 215–220. https://doi.org/10.1080/165019702760279206

Ardila, A., & Rosselli, M. (1989). Neuropsychological characteristics of normal aging. Developmental Neuropsychology, 5(4), 307–320. https://doi.org/10.1080/87565648909540441

Bellman, R. (1957). Dynamic programming. Princeton: Princeton University Press.

Binder, J., Marshall, R., Lazar, R., Benjamin, J., & Mohr, J. P. (1992). Distinct Syndromes of Hemineglect. Archives of Neurology, 49(11), 1187–1194. https://doi.org/10.1001/archneur.1992.00530350109026

Brucki, S. M. D., & Nitrini, R. (2008). Cancellation task in very low educated people. Archives of Clinical Neuropsychology. https://doi.org/10.1016/j.acn.2007.11.003

Buxbaum, L. J., Ferraro, M. K., Veramonti, T., Farne, A., Whyte, J., Ladavas, E., … Coslett, H. B. (2004). Hemispatial neglect: Subtypes, neuroanatomy, and disability. Neurology, 62(5), 749–756. https://doi.org/10.1212/01.WNL.0000113730.73031.F4

Byrd, D. E., Touradji, P., Tang, M.-X., & Manly, J. T. (2004). Cancellation test performance in African American, Hispanic, and White elderly. Journal of the International Neuropsychological Society, 10, 401–411.

Dalmaijer, E. S., Li, K. M. S., Gorgoraptis, N., Leff, A. P., Cohen, D. L., Parton, A., … Malhotra, P. A. (2018). Randomised, double-blind, placebo-controlled crossover study of single-dose guanfacine in unilateral neglect following stroke. Journal of Neurology, Neurosurgery & Psychiatry, jnnp-2017-317338. https://doi.org/10.1136/jnnp-2017-317338

Dalmaijer, E. S., Van der Stigchel, S., Nijboer, T. C. W., Cornelissen, T. H. W., & Husain, M. (2015). CancellationTools: All-in-one software for administration and analysis of cancellation tasks. Behavior Research Methods, 47(4), 1065–1075. https://doi.org/10.3758/s13428-014-0522-7

Davies, A. D. M., & Davies, D. B. (1975). The Effects of Noise and Time of Day upon Age Differences in Performance at Two Checking Tasks. Ergonomics, 18(3), 321–336. https://doi.org/10.1080/00140137508931465

Donnelly, N., Guest, R., Fairhurst, M., Potter, J., Deighton, A., & Patel, M. (1999). Developing algorithms to enhance the sensitivity of cancellation tests of visuospatial neglect. Behavior Research Methods, Instruments, & Computers, 31(4), 668–673.

Geldmacher, D. S. (1996). Effects of stimulus number and target-to-distractor ratio on the performance of random array letter cancellation tasks. Brain and Cognition, 32, 405–415.

Geldmacher, D. S. (1998). Stimulus characteristics determine processing approach on random array letter-cancellation tasks. Brain and Cognition, 36, 346–354.

Geldmacher, D. S., Fritsch, T., & Riedel, T. M. (2000). Effects of Stimulus Properties and Age on Random-Array Letter Cancellation Tasks. Aging, Neuropsychology, and Cognition, 7(3), 194–204. https://doi.org/10.1076/1382-5585(200009)7:3;1-Q;FT194

Hills, E. C., & Geldmacher, D. S. (1998). The effect of character and array type on visual spatial search quality following traumatic brain injury. Brain Injury, 12(1), 69–76. https://doi.org/10.1080/026990598122872

Holm, S. (1979). A simple sequentially rejective multiple test procedure. Scandinavian Journal of Statistics, 6(2), 65–70.

Husain, M., & Rorden, C. (2003). Non-spatially lateralized mechanisms in hemispatial neglect. Nature Reviews Neuroscience, 4(1), 26–36. https://doi.org/10.1038/nrn1005

Jain, A. K. (2010). Data clustering: 50 years beyond K-means. Pattern Recognition Letters, 31(8), 651–666. https://doi.org/10.1016/j.patrec.2009.09.011

Kaufman, L., & Rousseeuw, P. J. (Eds.). (1990). Finding Groups in Data. Hoboken, NJ, USA: John Wiley & Sons, Inc. https://doi.org/10.1002/9780470316801

Lowery, N., Ragland, J. D., Gur, R. C., Gur, R. E., & Moberg, P. J. (2004). Normative data for the symbol cancellation test in young healthy adults. Applied Neuropsychology, 11(4), 218–221.

Malhotra, P. A., Jager, H. R., Parton, A., Greenwood, R., Playford, E. D., Brown, M. M., & Driver, J. (2005). Spatial working memory capacity in unilateral neglect. Brain, 128(2), 424–435. https://doi.org/10.1093/brain/awh372

Malhotra, P. A., Parton, A. D., Greenwood, R., & Husain, M. (2006). Noradrenergic modulation of space exploration in visual neglect. Annals of Neurology, 59(1), 186–190. https://doi.org/10.1002/ana.20701

Mark, V. W., Woods, A. J., Ball, K. K., Roth, D. L., & Mennenmeier, M. (2004). Disorganized search on cancellation is not a consequence of neglect. Neurology, 63, 78–84.

Mazaux, J. M., Dartigues, J. F., Letenneur, L., Darriet, D., Wiart, L., Gagnon, M., … Boller, F. (1995). Visuo-spatial attention and psychomotor performance in elderly community residents: Effects of age, gender, and education. Journal of Clinical and Experimental Neuropsychology, 17(1), 71–81. https://doi.org/10.1080/13803399508406583

Mesulam, M.-M. (1985). Principles of Behavioural Neurology. Tests of directed attention and memory. Philadelphia, US: Davis.

Na, D. L., Adair, J. C., Kang, Y., Chung, C. S., Lee, K. H., & Heilman, K. M. (1999). Motor perseverative behavior on a line cancellation task. Neurology, 52(8), 1569–1569. https://doi.org/10.1212/WNL.52.8.1569

Nijboer, T. C. W., Kollen, B. J., & Kwakkel, G. (2013). Time course of visuospatial neglect early after stroke: A longitudinal cohort study. Cortex, 49(8), 2021–2027. https://doi.org/10.1016/j.cortex.2012.11.006

Nys, G. M. S., Van Zandvoort, M., Van der Worp, H., Kappelle, L. J., & De Haan, E. H. F. (2006). Neuropsychological and neuroanatomical correlates of perseverative responses in subacute stroke. Brain, 129(8), 2148–2157. https://doi.org/10.1093/brain/awl199

Parton, A. D., Malhotra, P. A., Nachev, P., Ames, D., Ball, J., Chataway, J., & Husain, M. (2006). Space re-exploration in hemispatial neglect. NeuroReport, 17(8), 833–836.

Rabuffetti, M., Farina, E., Alberoni, M., Pellegatta, D., Appollonio, I., Affanni, P., … Ferrarin, M. (2012). Spatio-Temporal Features of Visual Exploration in Unilaterally Brain-Damaged Subjects with or without Neglect: Results from a Touchscreen Test. PLoS ONE, 7(2), e31511. https://doi.org/10.1371/journal.pone.0031511

Rousseeuw, P. (1987). Silhouettes: A graphical aid to the interpretation and validation of cluster analysis. Journal of Computational and Applied Mathematics, 20, 53–65. https://doi.org/10.1016/0377-0427(87)90125-7

Rusconi, M. L., Maravita, A., Bottini, G., & Vallar, G. (2002). Is the intact side really intact? Perseverative responses in patients with unilateral neglect: a productive manifestation. Neuropsychologia, 40(6), 594–604. https://doi.org/10.1016/S0028-3932(01)00160-9

Samuelsson, H., Hjelmquist, E. K. E., Jensen, C., & Blomstrand, C. (2002). Search pattern in a verbally reported visual scanning test in patients showing spatial neglect. Journal of the International Neuropsychological Society, 8(03), 382–394. https://doi.org/10.1017/S1355617702813194

Saykin, A. J., Gur, R. C., Gur, R. E., Shtasel, D. L., Flannery, K. A., Mozley, L. H., … Mozley, P. D. (1995). Normative neuropsychological test performance: effects of age, education, gender, and ethnicity. Applied Neuropsychology, 2, 79–88.

Ten Brink, A. F., Biesbroek, M. J., Kuijf, H. J., Van der Stigchel, S., Oort, Q., Visser-Meily, J. M. A., & Nijboer, T. C. W. (2016). The right hemisphere is dominant in organization of visual search—A study in stroke patients. Behavioural Brain Research, 304, 71–79. https://doi.org/10.1016/j.bbr.2016.02.004

Ten Brink, A. F., Van der Stigchel, S., Visser-Meily, J. M. A., & Nijboer, T. C. W. (2015). You never know where you are going until you know where you have been: Disorganized search after stroke. Journal of Neuropsychology. https://doi.org/10.1111/jnp.12068

Ten Brink, A. F., Verwer, J. H., Biesbroek, J. M., Visser-Meily, J. M. A., & Nijboer, T. C. W. (2017). Differences between left- and right-sided neglect revisited: A large cohort study across multiple domains. Journal of Clinical and Experimental Neuropsychology, 39(7), 707–723. https://doi.org/10.1080/13803395.2016.1262333

Uttl, B., & Pilkenton-Taylor, C. (2001). Letter Cancellation Performance Across the Adult Life Span. The Clinical Neuropsychologist (Neuropsychology, Development and Cognition: Section D), 15(4), 521–530. https://doi.org/10.1076/clin.15.4.521.1881

Van der Maaten, L. J. P., & Hinton, G. E. (2008). Visualizing high-dimensional data using t-SNE. Journal of Machine Learning Research, 9, 2579–2605.

Warren, M., Moore, J. M., & Vogtle, L. K. (2008). Search Performance of Healthy Adults on Cancellation Tests. American Journal of Occupational Therapy, 62(5), 588–594. https://doi.org/10.5014/ajot.62.5.588

Weintraub, S., & Mesulam, M.-M. (1988). Visual hemispatial inattention: stimulus parameters and exploratory strategies. Journal of Neurology, Neurosurgery & Psychiatry, 51, 1481–1488.

Woods, A. J., Göksun, T., Chatterjee, A., Zelonis, S., Mehta, A., & Smith, S. E. (2013). The development of organized visual search. Acta Psychologica, 143(2), 191–199. https://doi.org/10.1016/j.actpsy.2013.03.008

Woods, A. J., & Mark, V. W. (2007). Convergent validity of executive organization measures on cancellation. Journal of Clinical and Experimental Neuropsychology, 29(7), 719–723. https://doi.org/10.1080/13825580600954264

Woods, A. J., Mark, V. W., Pitts, A. C., & Mennemeier, M. (2011). Pervasive Cognitive Impairment in Acute Rehabilitation Inpatients Without Brain Injury. PM&R, 3(5), 426–432. https://doi.org/10.1016/j.pmrj.2011.02.018

